# DNA-PKcs Inhibition Improves Sequential Gene Insertion of the Full-Length *CFTR* cDNA in Airway Stem Cells

**DOI:** 10.1101/2024.08.12.607571

**Authors:** Jacob T. Stack, Rachael E. Rayner, Reza Nouri, Carlos J. Suarez, Sun Hee Kim, Karen L. Kanke, Tatyana A. Vetter, Estelle Cormet-Boyaka, Sriram Vaidyanathan

**Author notes:** Correspondence: Sriram Vaidyanathan, PhD. Center for Gene Therapy, Abigail Wexner Research Institute, Nationwide Children’s Hospital, 575 Children’s Crossroad, Columbus, OH – 43215.

## Abstract

Cystic fibrosis (CF) is caused by mutations in the cystic fibrosis transmembrane conductance regulator (*CFTR*) gene. Although many people with CF (pwCF) are treated using CFTR modulators, some are non-responsive due to their genotype or other uncharacterized reasons. Autologous airway stem cell therapies, in which the *CFTR* cDNA has been replaced, may enable a durable therapy for all pwCF. Previously, CRISPR-Cas9 with two AAVs was used to sequentially insert two halves of the *CFTR* cDNA and an enrichment cassette into the *CFTR* locus. However, the editing efficiency was <10% and required enrichment to restore CFTR function. Further improvement in gene insertion may enhance cell therapy production. To improve *CFTR* cDNA insertion in human airway basal stem cells (ABCs), we evaluated the use of the small molecules AZD7648 and ART558 which inhibit non-homologous end joining (NHEJ) and micro-homology mediated end joining (MMEJ). Adding AZD7648 alone improved gene insertion by 2-3-fold. Adding both ART558 and AZD7648 improved gene insertion but induced toxicity. ABCs edited in the presence of AZD7648 produced differentiated airway epithelial sheets with restored CFTR function after enrichment. Adding AZD7648 did not increase off-target editing. Further studies are necessary to validate if AZD7648 treatment enriches cells with oncogenic mutations.

## INTRODUCTION

Cystic Fibrosis (CF) is a monogenic disorder that results from mutations in the cystic fibrosis transmembrane conductance regulator (*CFTR*) gene. CF is a fatal disease in childhood if left untreated.^1^ CFTR is a chloride/bicarbonate ion channel protein present in many epithelia, including the airways, intestines, pancreatic ducts and vas deferens.^2^ Lack of CFTR function causes the buildup of thick mucus with reduced bacterial killing ability which results in repeated lung infections and lung failure.^3–5^ Therapeutic interventions prior to 2012 did not restore CFTR function directly but instead focused on infection clearance, loosening mucus and improving nutrient absorption.^1^ Over the past decade, small molecules that restore the folding and ion channel function of mutant CFTR (CFTR modulators) have been developed.^6, 7^ CFTR modulators are effective in treating pwCF affected by a variety of pathogenic variants including the F508del variant seen in ∼90% of pwCF in Europe and North America.^8, 9^ Although CFTR modulators have dramatically improved outcomes for pwCF, ∼10% of pwCF with European ancestry and between 30-50% of pwCF from other ancestries cannot be treated with these drugs. ^10, 11^ In addition, modulators are not curative and must be taken daily for the life of a pwCF. Furthermore, modulators can cause severe side-effects including neurocognitive symptoms.^12^ Thus, there is a continued interest in other approaches to restore CFTR production and function which may enable a durable treatment for all pwCF.^10, 11^

Genetic therapies which deliver a functional copy of *CFTR* into airway cells have been proposed to treat CF lung disease and would enable treatment of all pwCF regardless of the causal mutations.^13^ Unfortunately, gene therapy trials using lipid nanoparticles (LNPs) and adeno-associated virus (AAV) vectors to transiently deliver *CFTR* mRNA or cDNA have been unsuccessful.^14–16^ Although newer LNPs, AAV6s and related viruses are being developed to improve transduction of airway cells in the presence of the thick mucus seen in pwCF, the approach will still require redosing.^17–21^ This can be particularly challenging with viral vectors because of anti-viral immunity mediated by the initial dose.^22, 23^ Lentiviral insertion of CFTR offers an alternative but it does not restore native gene expression and poses a small but documented risk for insertional mutagenesis.^24–27^ Genome editing to correct *CFTR* individual mutations or replace the *CFTR* cDNA has been proposed as an alternative approach to transient gene addition or lentiviral mediate gene insertion.^10, 28–31^ Correction of *CFTR* variants or replacement of the *CFTR* cDNA in airway basal stem cells (ABCs), which give rise to other cell types in the airway epithelium, may result in a durable therapy for CF.

CRISPR-Cas9 is commonly used to perform gene editing.^32–34^ In this method, Cas9 induces a double-stranded break in the target locus which is then repaired by several competing DNA repair pathways. Gene correction relies on the homologous recombination (HR) DNA repair pathway which requires the delivery of a template DNA (HR template) containing the correction sequence or *CFTR* cDNA. Delivery of the HR template using AAV results in highly efficient gene insertion in many therapeutically relevant primary stem cells *in vitro*, including ABCs.^35^ ^28, 36^ *In vivo* genome editing of ABCs to insert genes is limited by inefficient delivery of cargo into those cells and the absence of HR in quiescent ABCs *in vivo*.^34^ An *ex-vivo* cell therapy approach to correct CF causing mutations in ABCs *in vitro* followed by their transplantation has been proposed.^28^ Although it is possible to correct individual mutations with high efficiencies using HR-based gene correction as well as other editing approaches, the replacement of the *CFTR* cDNA is desirable due to the large number of variants scattered throughout the CFTR gene.^28–30^ One challenge with inserting the *CFTR* cDNA in ABCs using this approach is that a single AAV vector does not fit the *CFTR* cDNA and homology arms (HAs) needed for gene insertion. Hence, a sequential gene insertion approach was used to insert the entire *CFTR* cDNA into the endogenous locus.^10^ The study used two AAVs to insert the two halves of *CFTR* cDNA and an enrichment cassette expressing truncated CD19 (tCD19), into the *CFTR* locus and enabled the development of a universal CFTR restoration strategy applicable to almost all pwCF (universal strategy). It was necessary to use tCD19 as an enrichment tag because CFTR is not expressed in ABCs at high levels.^37^ Expression of tCD19 is mediated by the human phosphoglycerate kinase (PGK) promoter which is constitutive. The universal strategy yielded <10% tCD19^+^ cells and required the enrichment of tCD19^+^ cells using fluorescent or magnetic activated cell sorting (FACS or MACS).^10^ The enriched tCD19^+^ cells produced differentiated airway epithelial sheets with CFTR function comparable to that seen in non-CF controls. Furthermore, in a follow-up study it was shown that the strategy does not alter CFTR regulation and does not compromise the regenerative potential of edited ABCs.^38^ Despite this promising development, one drawback of the approach is that >90% of cultured cells are lost in the enrichment process which limits the cell yield to 1-10 million cells. This yield may be sufficient to replace the sinus epithelium and treat CF sinus disease.^28^

However, studies in mice have suggested that 60-100 million cells may be necessary to replace the lower airway epithelium in humans.^39–41^ Thus, further improvement in gene insertion efficiency will improve our ability to produce a gene edited airway stem cell therapy for treating CF lung disease using the universal strategy.

Studies have shown that the inhibition of non-homologous end joining (NHEJ) and micro-homology mediated end joining (MMEJ) which compete with HR improves gene insertion.^42–44^ DNA dependent protein kinase-catalytic subunit (DNA-PKcs) and DNA polymerase theta (PolΘ) are involved in NHEJ and MMEJ respectively. Previous studies have knocked out DNA-PKcs and PolΘ to increase gene insertion using HR.^44^ Other studies have increased HR using AAV and ssDNA templates by transiently inhibiting NHEJ and MMEJ using small molecules.^42, 45^ The use of the DNA PKcs inhibitor AZD7648 improved correction of the F508del mutation in airway basal stem cells by 2-3 fold.^42^ The combination of DNA-PKcs inhibitor to inhibit NHEJ and PolΘ inhibitor to inhibit MMEJ resulted in editing efficiencies of up to 80% in multiple primary stem cells using ssDNA templates.^43^ Despite these reports, it is unclear how the strategies impact sequential gene insertion which is used to insert the *CFTR* cDNA in the universal strategy. Moreover, the combined inhibition of DNA-PKcs and PolΘ was not tested in primary human airway cells.^45^ Furthermore, the impact of DNA-PKcs and PolΘ inhibition on the ability of ABCs to differentiate into airway cells and produce functional CFTR is still unknown.

Apart from improving gene insertion, it is important to characterize the safety of inhibiting DNA repair pathways. Off-target editing by Cas9 is a common safety concern that is specific to the sgRNA and Cas9 variant used to generate the edited cells.^46^ Our previous research has shown that the sgRNA used in the universal strategy does not cause significant off-target editing when used with high-fidelity Cas9.^10, 47^ Previous studies using DNA-PKcs inhibition have reported an increase in off-target edits but the frequency of alleles with off-target editing was still low (0.03-3% edited alleles).^42^ Another concern associated with genome editing is the enrichment of cells with oncogenic changes.^48, 49^ The use of Cas9 ribonuclear protein (RNP) for editing limits the window of genome editing to 2-4 days and thus has not resulted in an enrichment of cells with oncogenic changes when used to edit primary stem cells.^10, 50, 51^ However, the impact of inhibiting DNA repair pathways on the enrichment of cells with oncogenic changes needs further characterization.

In this study, we evaluated if the use of AZD7648, a DNA-PKcs inhibitor, alone or in combination with ART558, a PolΘ inhibitor, best improved gene insertion in ABCs and further assessed their impact on cell proliferation and differentiation. We evaluated the ability of both strategies to improve CFTR cDNA insertion using the universal strategy. Furthermore, we evaluated the restoration of CFTR function in differentiated airway cells derived from CF ABCs edited in the presence of AZD7648. To evaluate safety, we characterized the impact of DNA-PKcs inhibition using AZD7648 on off-target activity and the enrichment of cells with oncogenic changes.

## RESULTS

### Addition of AZD7648 improves gene insertion in primary human airway basal stem cells (ABCs)

We first tested our hypothesis by evaluating if the use of AZD7648 improves the insertion of a cassette expressing GFP under the control of an SFFV promoter in primary ABCs (Figure 1A). DSBs were introduced in the CFTR locus using Cas9 RNP and a previously reported sgRNA targeting exon 1 of CFTR.^10^ The template coding for GFP was delivered using AAV6 serotype 6 (AAV6) after the Cas9 RNP and chemically modified sgRNA were electroporated into the ABCs. Gene insertion in ABCs was assessed in the presence and absence of AZD7648. A representative flow cytometry plot shows that edited ABCs are GFP^+^ (Figure 1B). Without AZD7648, 34.8 ± 21.5% of edited ABCs from three different donors were positive for GFP. Inhibiting NHEJ through the addition of 0.5 µM AZD7648 significantly increased editing efficiency (p<0.05) to 61.3 ± 26.1% when assessed using Wilcoxon matched pairs significant rank test (Figure 1C). When compared to the vehicle control, 4-5 days after editing, AZD7648 did not reduce cell proliferation significantly (Figure 1D). We validated insertion of the GFP cassette in the CFTR locus by performing an IN-OUT polymerase chain reaction (PCR) in which one primer binds a part of the transgene cassette and the other primer binds a genomic sequence outside the arms of homology (Figure S1A). We observed a PCR product only in ABCs edited in the presence of AZD7648 but not in the controls (Figure S1B). Upon sequencing, the sequence of the amplicon matched the transgene cassette thus confirming targeted insertion in the CFTR locus (Figure S1C-D).

**Figure 1.**
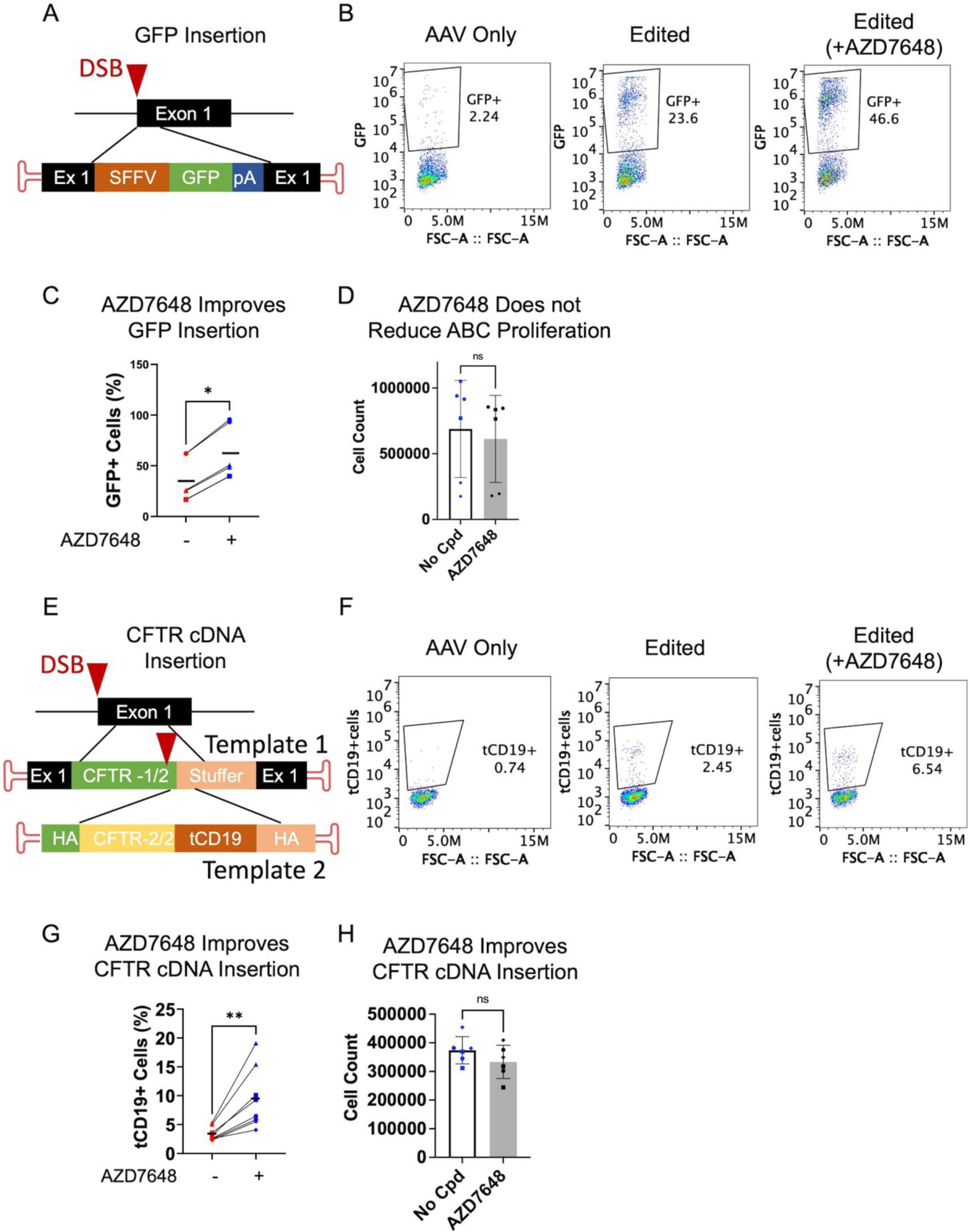
Insertion of GFP and *CFTR* cDNA in non-CF ABCs in the presence of AZD7648. (A) Schematic of the template used for knocking in a GFP expression cassette into exon 1 of *CFTR* using Cas9 RNP/AAV based gene editing. (B) Representative flow cytometry plots of cells edited in the presence of AZD7648 and AAV only controls. (C) Insertion of GFP in three biological replicates (2 technical replicates per donor). (D) Proliferation of ABCs edited to insert GFP in the presence of AZD7648 was not different from ABCs edited without AZD7648 measured 5 days after editing. (E) Schematic showing insertion of the *CFTR* cDNA and *tCD19* enrichment tag into exon 1 of the *CFTR* locus using the sequential insertion strategy (universal strategy). The second insertion is initiated by including the same sgRNA sequence from exon 1 of *CFTR* at the end of the first template. The *CFTR* cDNA is followed by a BGH polyA tail. The tCD19 cassette is driven by a PGK promoter and has an SV40 polyA tail. (F) Representative flow cytometry plots depicting tCD19 expression in the presence of AZD7648 or AAV only controls. The AAV only plot depicts the minimal episomal expression from the use of AAV. (G) Expression of tCD19 in edited non-CF ABCs in the presence of AZD7648. Presented data is from four biological replicates (2 technical replicates per donor). (H) Proliferation of ABCs after *CFTR* cDNA and *tCD19* insertion in the presence of AZD7648 in comparison to ABCs edited without AZD7648 measured on day 5 after editing. All statistical significance was assessed using Wilcoxon matched-pairs significant rank test. **, * represent p < 0.01 and 0.05 respectively. Cpd = compound.

Since recent studies have reported that DNA-PKcs inhibitors improve AAV transduction in differentiated airway cells, we evaluated if AZD7648 treatment improves gene insertion by improving AAV transduction.^52^ ABCs were treated with AAV alone in the presence or absence of AZD7648 without any Cas9. There was no difference in the percent of GFP^+^ ABCs between the two conditions when measured 24 h after transduction (Figure S1E).

Given the improvement in gene insertion using a single template, we then investigated if AZD7648 improves *CFTR* cDNA replacement using the universal strategy. The universal strategy uses the same sgRNA as the GFP insertion experiment but involves the sequential gene insertion of the *CFTR* cDNA and tCD19 expression cassette delivered using two different AAVs (Figure 1E). Editing efficiency was determined by characterizing the percent of tCD19^+^ cells using flow cytometry (Figure 1F). Without AZD7648, integration of the *CFTR* cDNA was 3.4 ± 1.2%. Inhibiting NHEJ through the addition of AZD7648 significantly increased editing to 9.5 ± 5.3% (p < 0.01) when assessed using Wilcoxon matched-pairs significant rank test (Figure 1G). Similar to the experiments inserting GFP, using AZD7648 to enhance gene insertion using the universal strategy did not reduce cell proliferation significantly (Figure 1H).

### Addition of ART558 with AZD7648 improves gene insertion further but causes cell death

Although the use of AZD7648 increased *CFTR* cDNA insertion using the universal strategy, <10% tCD19^+^ cells were obtained on average. Therefore, we investigated if combining AZD7648 with ART558, which is reported to inhibit PolΘ, further improves gene insertion. Previous studies have reported that the simultaneous use of small molecules inhibitors of DNA-PKcs and PolΘ results in improved gene insertion when compared to the use of a DNA-PKcs inhibitor alone.^43, 53^ The use of ART558 alone did not improve gene insertion when assessed using our previously reported F508del gene correction system (Figure S2A-B).^28^ We then added AZD7648 and ART558 while editing primary ABCs using Cas9/sgRNA and a single AAV template expressing GFP. Representative plots show GFP^+^ cells measured using flow cytometry (Figure 2A). With no AZD7648 or ART558 compounds, 23 ± 18% were GFP^+^ (Figure 2B). Inhibiting NHEJ with AZD7648 increased editing to 57 ± 23% GFP^+^ cells and inhibiting NHEJ and MMEJ using both AZD7648 and ART558 resulted in 72 ± 21% GFP^+^ cells. When assessed with the Wilcoxon matched-pairs significant rank test these differences were statistically significant (p < 0.001 and p < 0.05, respectively) (Figure 2B). However, treatment with both ART558 and AZD7648 reduced cell proliferation to 55% of ABCs edited without any compound compared. (Figure 2C). This reduction in cell proliferation was statistically significant when assessed using the ratio paired T-test (p < 0.05).

**Figure 2.**
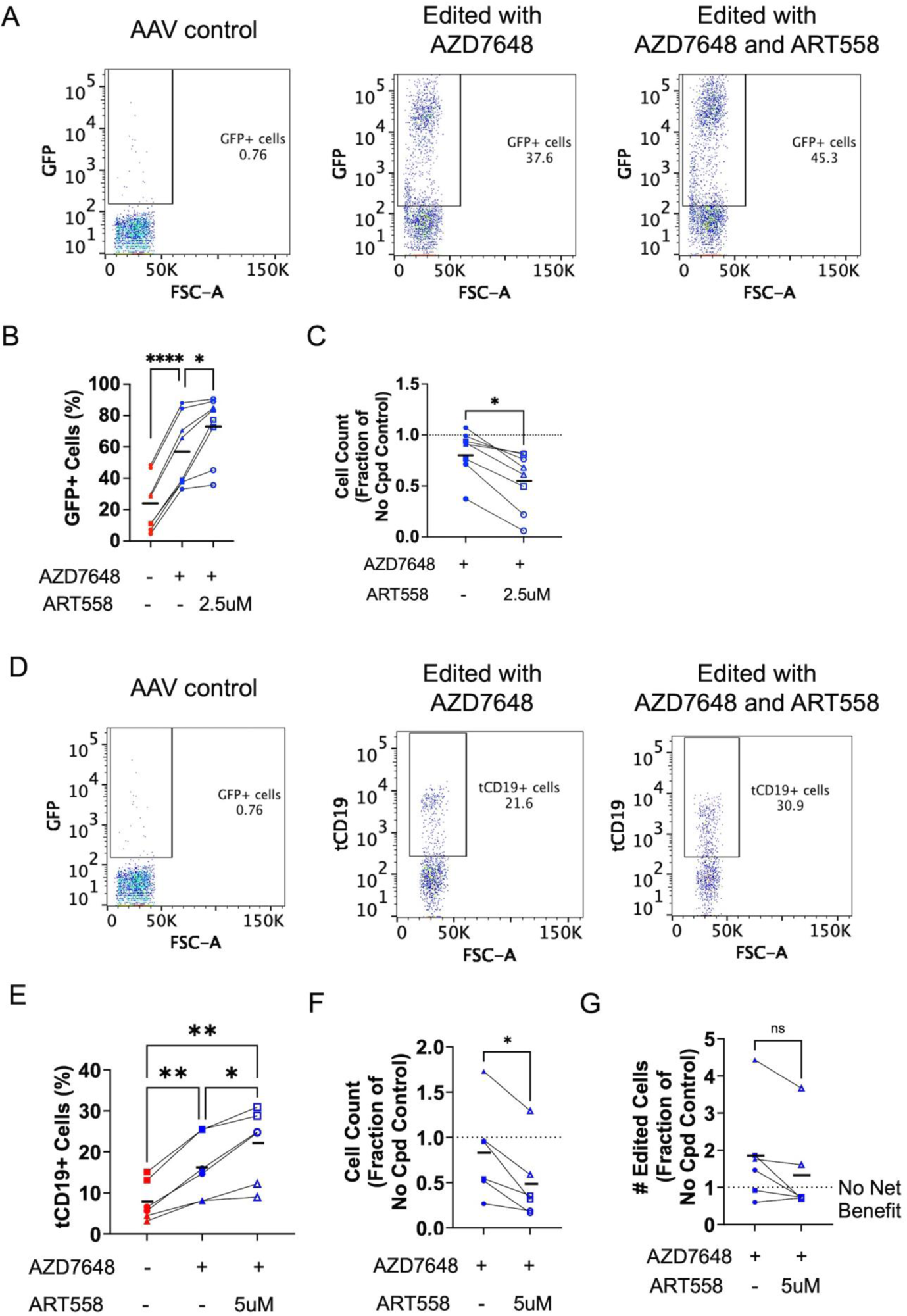
Insertion of GFP and tCD19 in non-CF ABCs in the presence of ART558 and AZD7648. (A) Representative flow cytometry plot showing the percent of GFP^+^ cells after genome editing performed in the presence of AZD7648 alone, or AZD7648 and ART558. (B) GFP^+^ cells measured from three different donors with the addition of AZD7648 alone, or AZD7648 and ART558. (C) Cell proliferation assessed when AZD7648 and ART558 compared with the AZD7648 only condition. (D) Representative flow cytometry plots of edited non-CF ABCs edited using the universal strategy in the presence of AZD7648 alone, or AZD7648 and ART558. (E) tCD19 expression in non-CF ABCs with the addition of AZD7648 alone, or AZD7648 and ART558. (F) Cell proliferation of ABCs edited using the universal strategy in the presence of AZD7648 only, or AZD7648 and ART558. (G) Relative edited cell yield between the AZD7648 only, or AZD7648 and ART558 conditions. All statistical significance was assessed using one-way ANOVA followed by Tukey’s test, or by ratio paired T-test. ****, **, * represent p < 0.0005, 0.01and 0.05 respectively. Cpd = compound.

AZD7648 and ART558 were added while editing ABCs using the universal strategy to determine if the inhibition of NHEJ and MMEJ affects sequential gene insertion differently. Representative plots show tCD19^+^ cells measured using flow cytometry (Figure 2D). 8.0 ± 4.9% of ABCs edited without AZD7648 or ART558 were tCD19^+^. Inhibiting NHEJ increased tCD19^+^ ABCs to 16.3 ± 7.8% and inhibiting both NHEJ and MMEJ yielded 21.8 ± 9.0% tCD19^+^ cells (Figure 2E). The addition of AZD7648 significantly increased editing by the universal strategy (p < 0.01) when assessed using ANOVA followed by Tukey’s multiple comparisons test. Unlike the experiments inserting GFP, the addition of ART558 only showed an increase in gene insertion at 5µM and showed no improvement in CFTR cDNA insertion at 2.5µM or 1µM (Figure S2C). ART558 with AZD7648 also significantly increased editing (p < 0.05) beyond the AZD7648 only condition when assessed with the Tukey’s multiple comparisons test. However, similar to the single AAV template, cell proliferation was reduced by ∼51% with the addition of both AZD7648 and ART558 (Figure 2F) and this difference was statistically significant (p < 0.01) when assessed using a ratio paired T-test. Lower concentrations of ART558 which did not improve gene insertion showed reduced toxicity (Figure S2D). Overall, the increase in integration of *CFTR* using both AZD7648 and ART558 is offset by the reduction in cell proliferation. To determine if using a different PolΘ inhibitor could improve cell yield, we tested RP-6685 and ART812. However, we observed no significant increase in editing efficiency compared to ART558 (Figure S2E). Overall, the addition of ART558 increased *CFTR* cDNA insertion by ∼1.3 times, but also reduced cell proliferation by ∼1.7 times when compared to the AZD7648 only condition. We attempted to quantify the trade-off between improved *CFTR* cDNA insertion and reduced viability. We multiplied the total number of cells by the percent of cells modified to obtain the total number of tCD19^+^ cells for each condition. We divided the number of tCD19^+^ cells obtained in the presence of the compounds by the number of tCD19^+^ cells obtained without compounds to obtain the relative cell yield. The addition of ART558 did not improve overall cell yield significantly when compared to the addition of AZD7648 alone (Figure 2G). We evaluated if the different compounds triggered cell death programs by staining for Annexin V and propidium iodide (PI). Treatment with ART558 and AZD7648 did not result in an increase in ABCs positive for Annexin V alone or both Annexin V and PI (Figure S3).

### Insertion of the full-length *CFTR* cDNA and enrichment in primary ABCs obtained from donors with CF

We then tested the universal strategy with AZD7648 on CF donor ABCs. Similar to the non-CF donors, adding AZD7648 to edited cells improved gene insertion by ∼2-fold. Insertion in the no compound condition was 11.0 ± 6.4% while inhibiting NHEJ with AZD7648 significantly increased editing to 21.0 ± 15.9% when assessed using a paired t-test (p < 0.05) (Figure 3A). The number of cells counted on day 5 after editing was not significantly different between ABCs edited in the presence or absence of AZD7648 (Figure 3B). Significance was determined by paired T-test. After enrichment of tCD19^+^ cells using MACS, populations with >50% tCD19^+^ cells were obtained (Figure 3C). Basal cell markers, P63 and cytokeratin 5 (KRT5), were evaluated via flow cytometry (Figure 3D). The expression of P63 and KRT5 in edited CF donor cells treated with no compound and with AZD7648 were not significantly different from the expression of these basal cell markers in unedited CF basal cells when assessed using one way ANOVA (Figure 3D).

**Figure 3.**
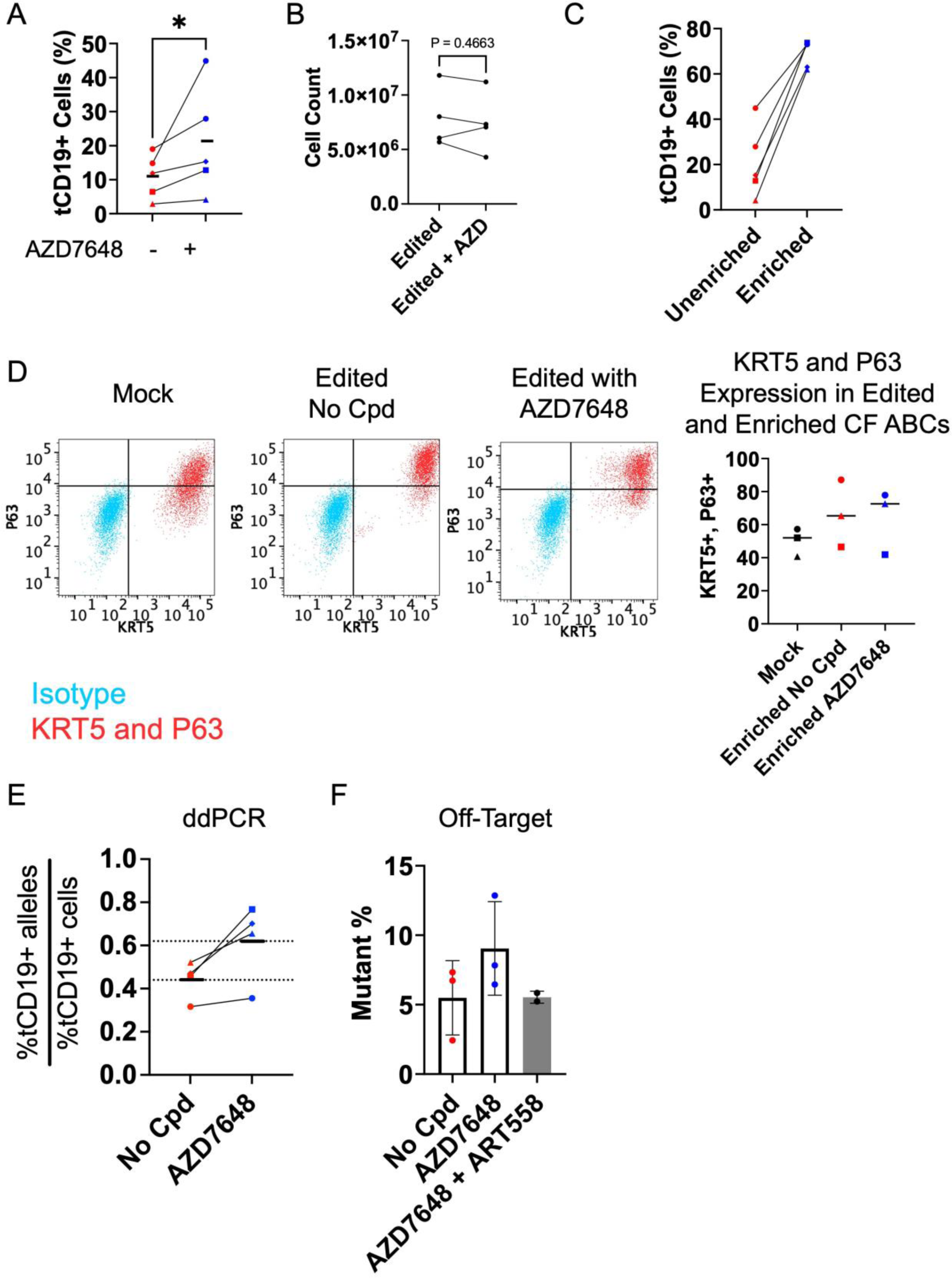
Primary human CF ABCs edited for *CFTR* cDNA insertion in the presence of AZD7648. (A) Insertion of tCD19 in CF-donor ABCs edited using the universal strategy. (B) The proliferation of edited CF ABCs was assessed 5 days after editing, with or without AZD7648. (C) The edited cell population was enriched for tCD19^+^ cells. (D) The enriched population was stained and evaluated for the presence of basal stem cell markets, P63 and KRT5, via flow cytometry. There was no significant difference in the percent of cells positive for P63 and KRT5 between the different conditions tested. (E) CF ABCs edited with or without AZD7648 were assessed for allelic frequency of tCD19^+^ alleles. (F) Enriched, edited CF cells were sequenced for off-target activity associated with the one known off-target site for the sgRNA used in editing. Off-target editing was not significantly different between the different conditions by one way ANOVA. Cpd = compound. * represents p < 0.05 in all panels.

We evaluated the allelic editing frequency associated with using the universal strategy in the presence and absence of AZD7648. Droplet digital PCR (ddPCR) was used to determine the percent of alleles with *tCD19* relative to a reference genomic sequence. A ratio of the percent of *tCD19*^+^ alleles relative to the percent of tCD19^+^ cells was calculated. A ratio equal to 1 suggests both alleles were edited in the edited cell population, whereas a ratio of 0.5 indicates 1 of 2 alleles was edited. For samples corrected without AZD7648, the average ratio was 0.44 ± 0.09 and for samples corrected with AZD7648, the average ratio was 0.62 ± 0.18 (Figure 3E). Thus, AZD7648 also improves the fraction of cells with biallelic insertion of the *CFTR* cDNA.

### Impact of DNA-PKcs inhibition on the off-target activity of the sgRNA

The sgRNA used to guide Cas9 to exon 1 of the *CFTR* locus can also direct the Cas9 to off-target (OT) sites as Cas9 can tolerate mismatches between the sgRNA and gDNA.^10^ To reduce OT activity, we used high-fidelity Cas9 (HiFi Cas9).^47^ AZD7648 has been shown to increase OT insertions and deletions (INDELs) in other studies.^42^ Therefore, we characterized INDELs in a previously identified OT site associated with our sgRNA in chromosome 5 in ABCs from three different donors using next-generation sequencing (NGS).^10^ In the ABCs edited with no compound, OT INDELs occurred in 5.5 ± 2.7% of cells. The addition of AZD7648 increased OT activity to 9.1 ± 3.4%. The increase was not significant when assessed using one-way ANOVA. However, when cells were treated with both AZD7648 and ART558, OT activity did not increase (5.5 ± 0.4%) relative to the controls edited without either compound (Figure 3F).

### Restoration of CFTR function in CF donor samples

After affirming that the CF donor cells edited with AZD7648 and enriched retained KRT5 and P63 expression, we evaluated the restoration of CFTR function in fully differentiated airway epithelial sheets derived from edited ABCs. The enriched population of edited CF ABCs were cultured and differentiated at air-liquid interface (ALI). Differentiated epithelial sheets showed transepithelial resistances (TEER) comparable to unedited controls thus indicating that AZD7648 did not compromise differentiation (Figure 4A). We further confirmed differentiation using immunofluorescence to ascertain the presence of ciliated cells and secretory cells marked by acetylated tubulin and CD66 respectively (Figure 4B).^54^ We assessed mature CFTR production using immunoblotting. While immunoblotting, mature CFTR that has undergone all the necessary posttranslational modifications appears at a higher molecular weight (∼170kDa) which is often referred as “band C”. Immature CFTR which appears at a lower molecular weight (∼150 kDa) is referred as “band B”. Differentiated airway cells from unedited non-CF and CF samples edited with and without AZD7648, respectively, showed a mature CFTR band (lanes 1, 3, and 4). By way of contrast, the unedited control CF sample showed no mature CFTR band (lane 2, Figure 4C).

**Figure 4.**
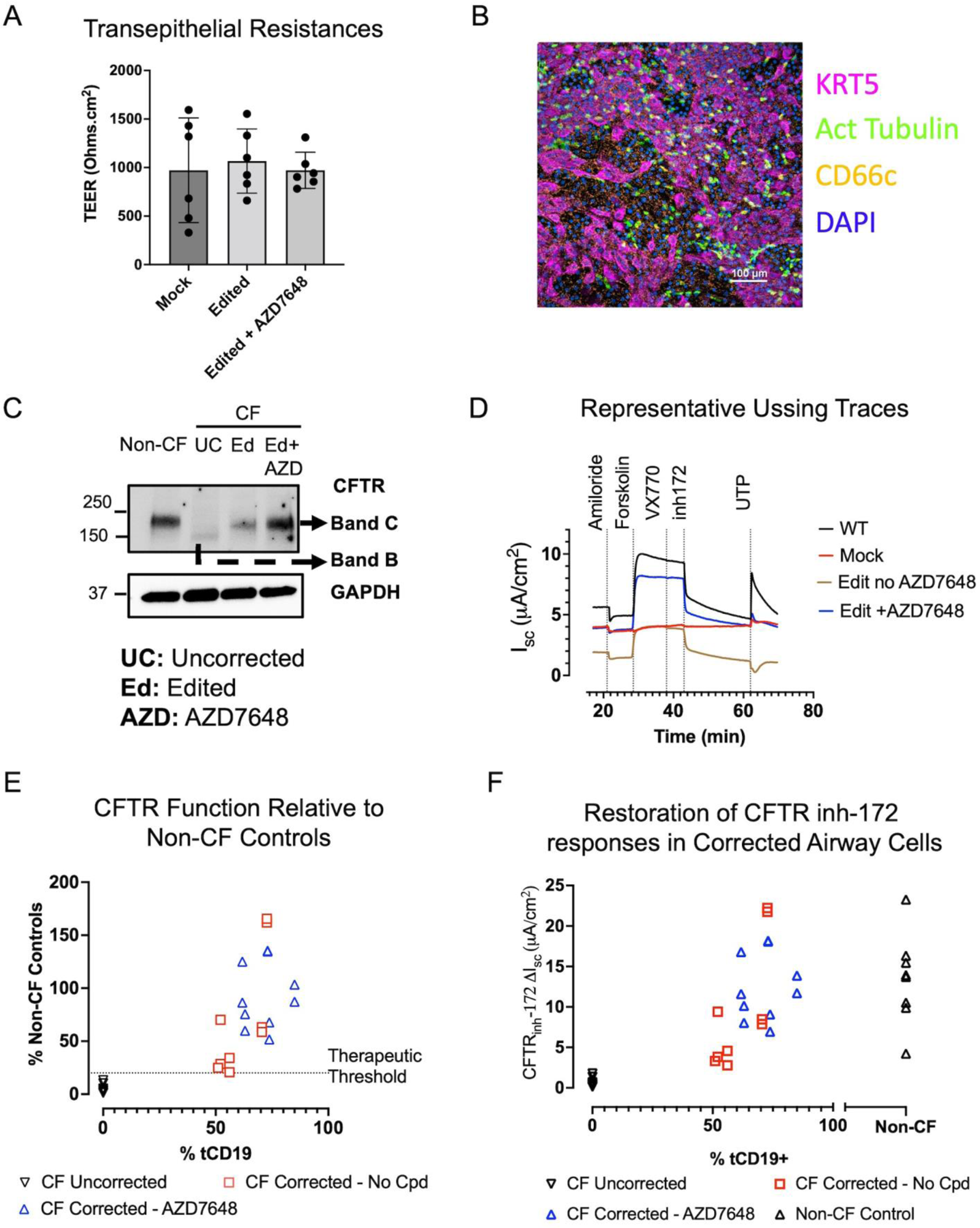
Differentiation and expression of CFTR in edited CF airway cells after differentiation on ALI transwells. WT non-CF ABCs and edited CF ABCS (with or without AZD7648) were differentiated on ALI transwells for at least 4 weeks. (A) After differentiation, transepithelial electrical resistance (TEER) was assessed in unedited CF controls (mock) and edited (No AZD7648 and AZD7648) samples. (B) Differentiated ALI cultures contained ciliated cells expressing acetylated tubulin and secretory cells positive for CD66. (C)CFTR protein expression in differentiated airway cells from non-CF, uncorrected CF and corrected CF samples (with and without AZD7648) was assessed using immunoblotting. (D) Ussing chamber analysis was conducted to evaluate CFTR function of differentiated ALI cultures. Representative traces from fully differentiated ALI cultures derived from non-CF ABCs (WT), unedited CF ABCs (Mock), CF ABCs edited without AZD7648, and CF ABCs edited with AZD7648 are displayed. (E) Percent of CFTR function (assessed from CFTR_inh_-172 response) in CF edited and unedited samples relative to non-CF controls is plotted against %tCD19*^+^* cells. 20% CFTR function relative to non-CF controls is thought to be therapeutically relevant. (F) Change (ΔIsc) in CFTR inhibition as assessed from CFTR_inh_-172 responses. When compared using one way ANOVA followed by Tukey’s test, the average ΔIsc values from CF samples corrected with or without AZD7648 were significantly higher than ΔIsc values uncorrected CF samples (p < 0.005). They were not significantly different from each other or the non-CF controls.

CFTR function was evaluated via Ussing chamber analysis. In differentiated non-CF airway cells (wild-type; WT), we observed an increase in short-circuit current upon the addition of forskolin (FSK) and a decrease in short-circuit current when CFTR inhibitor-172 (CFTR_inh_-172) was added. There was no change in CFTR current upon the addition of FSK or CFTR_inh_-172 in the unedited CF airway cells. The edited CF donor cells display similar current traces to non-CF cells upon the addition of these compounds (Figure 4D). The average change in short-circuit current after the addition of CFTR_inh_-172 was 13.4 ± 5.2 for non-CF, 0.6 ± 0.6 for uncorrected CF, 9.35 ± 7.5 for corrected CF samples without AZD7648, and 12.44 ± 4.1 for corrected CF samples with AZD7648. Upon analysis using a one-way ANOVA and Tukey’s post-hoc test, the change in CFTR_inh_-172 responsive current in the corrected CF samples with (p < 0.0001) and without (p = 0.0032) AZD7648 were significantly higher than the CFTR_inh_-172 responsive current observed in the uncorrected CF samples. Edited and enriched tCD19^+^ cells show CFTR function comparable to non-CF controls (Figure 4D-F). The restoration of CFTR function relative to the percent of tCD19^+^ cells is presented in Figure 4D-F. Table 1 lists the percent corrected cells and CFTR_inh_-172 response in each sample.

**Table 1:** Summary of percent editing in CF ABCs and relative restoration of CFTR function in differentiated airway cells derived from edited ABCs as measured by responses to CFTR_inh_-172.

| **No compounds** |  |  |  |
| --- | --- | --- | --- |
| Patient | Editing Efficiency<br>(% tCD19<sup>+</sup> Cells) | CFTR<sub>inh</sub>-172 currents<br>($\mu\text{A}/\text{cm}^2$) | Percentage of non-<br>CF-inhibitable current<br>(%) |
| 1 | 72.70 | -21.74 | 162.01 |
| 1 | 72.70 | -22.22 | 165.59 |
| 2 | 56.10 | -4.57 | 34.06 |
| 2 | 56.10 | -2.78 | 20.72 |
| 3 | 70.50 | -8.47 | 63.12 |
| 3 | 70.50 | -7.87 | 58.65 |
| 4 | 52.10 | -9.41 | 70.12 |
| 4 | 52.10 | -3.83 | 28.54 |
| 5 | 51.00 | -3.33 | 24.82 |
| **Average** | **61.53** | **-9.35 ± 7.54** | **69.74%** |
| **DNA-PKcs Inhibitor** |  |  |  |
| Patient | Editing Efficiency<br>(% tCD19<sup>+</sup> Cells) | CFTR<sub>inh</sub>-172 currents<br>($\mu\text{A}/\text{cm}^2$) | Percentage of non-<br>CF-inhibitable current<br>(%) |
| 1 | 73.00 | -18.08 | 134.73 |
| 1 | 73.00 | -18.17 | 135.41 |
| 2 | 73.80 | -9.08 | 67.67 |
| 2 | 73.80 | -6.94 | 51.72 |
| 3 | 85.00 | -13.87 | 103.36 |
| 3 | 85.00 | -11.71 | 87.26 |
| 4 | 61.80 | -16.78 | 125.05 |
| 4 | 61.80 | -11.59 | 86.37 |
| 5 | 63.00 | -8.03 | 59.84 |
| 5 | 63.00 | -10.13 | 75.49 |
| **Average** | **71.32** | **-12.44 ± 4.12** | **92.69%** |
| **Non-CF average: 13.4 ± 5.5$\mu\text{A}/\text{cm}^2$** |  |  |  |

### Impact of DNA-PKcs inhibition on enrichment of cells with oncogenic mutations

We used a next generation sequencing (NGS) panel evaluating 130 mutations associated with solid tumors to determine if any tumor causing genes were enriched in primary ABCs edited and enriched using the universal strategy in the presence of AZD7648. The results are presented in Table 2. There was no enrichment of oncogenic mutations in the two edited samples that were not exposed to AZD7648. In cells from the same two donors exposed to AZD7648, we observed no increased prevalence of oncogenic mutations in one donor. However, cells from the second donor showed an increase in cells containing a mutation in the DDX3X gene.

**Table 2:**
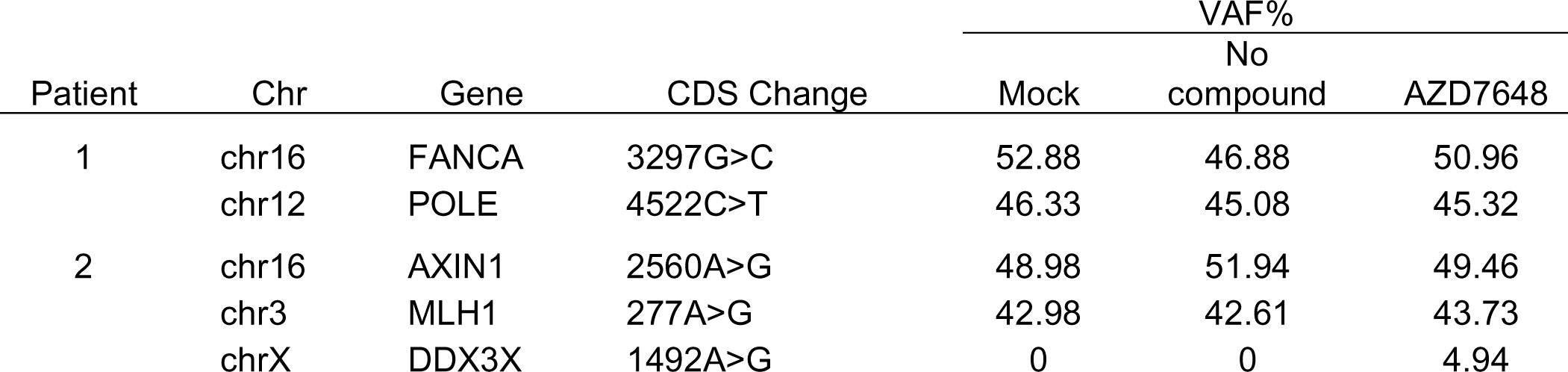
Summary of genes from STAMP panel with VAF > 5%.

The variable allele frequency of the oncogenic mutation in DDX3X increased from 0% in cells not subject to AZD7648 to 5% in cells treated with AZD7648.

## DISCUSSION

Recent studies have demonstrated that inhibition of DNA-PKcs and PolΘ improves gene insertion.^42, 45, 55^ The use of AZD7648 to inhibit DNA-PKcs improved gene insertion using AAV in multiple loci in multiple cell types.^42^ Since these studies focused on gene insertion of one cassette driven by a single HR reaction, the impact of DNA-PKcs inhibition on the sequential insertion of two templates was unknown. Here, we demonstrate that inhibition of DNA-PKcs also improves sequential gene insertion of the *CFTR* cDNA and tCD19 cassette in primary human ABCs by 2-3 fold. Previous studies have demonstrated that gene insertion using this two AAV strategy is sequential.^56^ There is variation between donors in the efficiency of *CFTR* cDNA insertion with and without AZD7648. However, the improvement in gene insertion using AZD7648 is consistently between 2-3 fold relative to the no compound controls from the same donor (Table S1).

In addition to AZD7648, studies have also reported that the combined inhibition of both DNA-PKcs and PolΘ improved gene correction of small segments (<50 bp) using ssDNA templates by 2-fold.^43^ Our results are consistent with these previous reports since we observe improved gene insertion of the GFP cassette and improved sequential insertion of the *CFTR* cDNA and *tCD19* cassette using both AZD7648 and ART558. However, the use of AZD7648 and ART558 to inhibit DNA-PKcs and PolΘ improves *CFTR* cDNA insertion using the two AAV strategy by only ∼40% and by 2-fold. Despite the improvement in gene insertion with the addition of ART558, there was a significant decrease in cell proliferation compared with the AZD7648 only condition. Although the cytotoxicity could be reduced by reducing ART558 concentration to 1µM, it also reduced gene insertion levels. It is possible that inhibition of NHEJ and MMEJ disrupts the ability of the cells to maintain genomic integrity and thus renders them non-viable. Although we did not measure genomic integrity, decreased off-target editing with the combined inhibition of DNA-PKcs and PolΘ is consistent with the hypothesis that inhibition of both DNA-PKcs and PolΘ favors the survival of cells that are able to repair the DSB successfully. Overall, there is no benefit of adding ART558 when the improvement in cell-yield is considered.

Although the use of AZD7648 improved the insertion of the CFTR cDNA by 2-3-fold, <20% ABCs were corrected on average. In our prior study, we observed that correction of <20% of alleles was insufficient to restore CFTR function.^28^ Therefore, enrichment remains necessary to produce a cell therapy with a therapeutically relevant percent of corrected cells. However, this 2-fold increase in gene insertion due of the use of AZD7648 will improve the yield of cells corrected using the universal strategy which can be useful for future cell therapy applications. In addition to improved cell yield, the cells edited using AZD7648 showed increased CFTR function (93% of non-CF controls) compared to the cells edited without AZD7648 (75% of non-CF controls) despite having a similar fraction of tCD19^+^ cells after enrichment. This is likely due to the fact that the use of AZD7648 increased the number of cells with bi-allelic correction as indicated by an increase in fraction of edited alleles as measured by ddPCR (Figure 3E). Thus, even though the use of AZD7648 does not enable us to avoid enrichment, it increases the level of CFTR restoration at least *in vitro*.

One of the limitations of our study is that we did not evaluate the impact of AZD7648 on the ability of ABCs to produce all the differentiated airway cells seen in an epithelium. Our results show that ABCs differentiate to produce a fully mature airway epithelium with ciliated and secretory cells. Our recent study investigated the regenerative potential of ABCs edited using the universal strategy by evaluating the production of different cell types through single-cell RNA sequencing.^38^ Similar studies may be needed to validate that AZD7648 does not alter the regenerative potential of edited ABCs.

Induced pluripotent stem cells (iPSCs) present an alternative source of cells for a cell therapy. Basal cells have been derived from iPSCs (iBasal cells) and have been shown to be able to produce a differentiated airway epithelium *in vivo*.^57, 58^ DNA-PKcs inhibition has been validated to improve gene insertion in iPSCs.^42, 43^ Therefore, the methods described in this study can be used to create genome edited iBasal cells.

In addition to evaluating the use of AZD7648 to improve gene insertion using the universal strategy, we also evaluated its impact on safety. NGS was used to evaluate off-target editing at the one known off-target site for the sgRNA used to insert the *CFTR* cDNA into the endogenous locus.^10^ The off-target site corresponds to an intergenic region in chromosome 5. This off-target site is 10 kb away from the closest gene (sideroflexin 1).^10^ The increase in off-target editing between the no compound control and the AZD7648 condition is not significant. Furthermore, the gene closest to this off-target locus (sideroflexin) is not active in airway cells.^10^ Further studies may be needed to assess changes in other possible off-target sites that were previously reported but did not show significant INDELs in the absence of AZD7648. Notably, we used a high-fidelity version of Cas9 in these studies that reduces off-target activity associated with this sgRNA by almost 10-20 fold.^10^ The use of wild-type Cas9 with AZD7648 may thus produce different results.

It has been recently reported that DNA-PKcs may play a role in limiting erroneous chromosomal rearrangements.^59^ Our recent study reported that the universal strategy results in translocations with the off-target site in <0.1% of alleles when measured using chromosomal aberrations analysis by single targeted linker-mediated PCR sequencing (CAST-seq).^38^ Notably, the only translocation that was reproducible between samples was mediated by off-target editing at the previously reported off-target site in chromosome 5.^10, 38^ The use of high fidelity Cas9 to limit activity in this off-target site is therefore even more necessary when using DNA-PKcs inhibition. We did not characterize translocations in cells treated with AZD7648. We posit that there may not be a significant increase in translocations because a translocation in exon 1 of *CFTR* would most likely abolish the CFTR function by disrupting the native *CFTR* promoter. Our data indicate that the airway cells edited in the presence of AZD7648 showed improved CFTR function relative to those edited in the absence of AZD7648. Nevertheless, further studies are necessary to evaluate if the use of AZD7648 during gene insertion increases the frequency of translocations prior to use in the production of a cell therapy for CF using this approach. In addition, further studies are necessary to evaluate if ABCs edited in the presence of AZD7648 maintain their regenerative potential in vivo.

We also used an NGS panel to evaluate the enrichment of cells with oncogenic mutations in 130 genes associated with solid tumors. A previous study had reported that the universal strategy does not result in an enrichment of cells with known oncogenic mutations.^10^ We observed no increase in the allelic frequencies of oncogenic mutations relative to the unedited controls in one of the two donors tested. However, we did observe an increase in cells with mutations in DDX3X only in the presence of AZD7648 in one donor. Overexpression of DDX3X is present in instances of head and neck squamous cell carcinoma and lung cancer.^60^ The mutation prevalence is within the 5% error range of the assay and only present in one of the two donors evaluated. Thus, it may be necessary to limit the exposure of cells to AZD7648 only to the duration needed for efficient HR (<24hrs). Furthermore, a test for the enrichment of oncogenic mutations may be necessary as a quality control step in the production of cell therapies if AZD7648 is used.

### Conclusions

Our experiments show that the inhibition of DNA-PKcs when inserting the full *CFTR* cDNA into the endogenous locus is most effective at improving edited cell yield. The addition of AZD7648 did not impact the cells’ ability to differentiate into functional airway epithelial sheets. The differentiated epithelium composed of cells edited with AZD7648 showed improved CFTR function when compared with the no compound control. The use of AZD7648 did not significantly increase off-target editing, but further studies need to be conducted to validate whether DNA-PKcs inhibition increases mutations in oncogenic genes.

## MATERIALS AND METHODS

### Subject details

De-identified primary airway epithelial basal cells (ABCs) were obtained from the Cure CF Columbus (C3) Epithelial Cell Core at Nationwide Children’s Hospital, Columbus OH.

### Method Details

#### Tissue culture of primary airway cells

Primary human ABCs were cultured in PneumaCult-Ex Plus (STEMCELL Technologies; 05040) and 10µM ROCK inhibitor (Y-27632; MedChemExpress; HY-10583) at 3,000 – 10,000 cells/cm^2^ in tissue-culture flasks. iMatrix (iMatrix-511 silk; Nacalai USA; 892021) was added to the cell culture media at time of plating. The volume of iMatrix to be added was calculated by halving the surface area of the plate and adding that value in microliters with the media.

#### Gene editing of ABCs

Primary ABCs were edited with sgRNA and HR templates specific to the universal correction method. The genomic target sequence for the sgRNA was TTCCAGAGGCGACCTCTGCA. The cells were cultured in PneumaCult-Ex Plus with 10 µM ROCK inhibitor and iMatrix. After 5 days of culture, TrypLE Express Enzyme (TrypLE) (Life Technologies; 12605036) was used to detach cells from the plates. The cells were resuspended in Opti-MEM (Life Technologies; 31985088) at a density of 10 million cells/mL. For electroporation, 20µL of cells were used for each condition (∼200,000 cells). Lonza 4D-Nucleofector (Lonza; AAF-1003B; AAF-1003X) was used for electroporation (nucleofection). HiFi Cas9 (6 µg; Integrated DNA Technologies, Coralville, IA, USA; 10007803) and 3.6 µg of 2’-O-methyl (M), 2’-O-methyl 3’phophorothioate modified sgRNA (MS-sgRNA) (Synthego Inc, Redwood City, CA, USA) (molar ratio = 1:2.5) were complexed at room temperature for 10min, mixed with the 20µL of cells suspended in Opti-MEM, and transferred to the nucleofector strip. Cells were then electroporated using the CA137 program and P3 solution. After electroporation, 80µL of Opti-MEM was added to each well. HR templates were delivered using AAV serotype 6. The HR sequences were packaged into AAV6 through commercial vendors (Vigene Inc or Signagen). While editing cells using the universal strategy, two AAVs carrying two-halves of the *CFTR* cDNA and tCD19 enrichment cassette were added at a multiplicity of infection (MOI) of 10^5^ genomes per cell, as determined by previous studies.^28^ While inserting GFP, the AAV carrying the HR template was added at an MOI of 10^5^ genomes per cell. Edited cells were transferred into multiple wells in a 12-well plate, 6-well plate, or T-75 flask such that the cell density was ∼5,000 cells/cm^2^. PneumaCult-Ex Plus, rock inhibitor, and iMatrix were added to each well/flask. A final concentration of 0.5µM AZD7648 (MedChem Express; HY-111783) and 5µM of ART558 (MedChem Express; HY-141520) were added to the cell-culture media. The media was replaced 24 hours after electroporation.

Primary ABCs were expanded and passaged for 4-5 days after editing and then replated to reduce episomal AAV expression and further expand edited cells for enrichment. Passaged ABCs were plated on tissue-culture treated flasks with a surface area of 175 cm^2^. Fifteen mL of PneumaCult-Ex Plus, 10µM ROCK inhibitor, and iMatrix were added to each flask. ABCs were detached from the plate with TrypLE 4-5 days after passaging and enriched for tCD19^+^ cells using MACS. tCD19 was detected using a mouse-anti-human CD19 antibody (Biolegend; 302256).

#### Enrichment via magnetic bead separation

ABCs were trypsinized using TryPLE and resuspended into MACS buffer composed of PBS with 2% bovine growth serum and 2 mM EDTA. The cells were enriched for tCD19^+^ cells with CD19 microbeads (Miltenyi Biotec; 130-050-301) using MACS LS columns (Miltenyi Biotec; 130-042-401) according to the manufacturers protocol. Prior to magnetic separation the cells were passed through a 5mL, 35µM Cell strainer snap cap (Fisher Scientific; 08-771-23) to obtain a single cell suspension to prevent the column from clogging.

#### Measuring insertion with flow cytometry

Cells were expanded after passaging or MACS for 4-5 days. The cells were collected using TryPLE and stained with FITC-anti-CD19 (Biolegend; 302256), AF647-anti-KRT5 (Abcam; ab193895), and anti-P63 (Biolegend; 687202). Isotype antibodies AF647 Rabbit IgG (Abcam; ab199093) for KRT5 and Mouse IgG2b (Biolegend; 401202) for P63 were used. The P63 antibody and isotype were unconjugated so PE goat-anti-mouse IgG (Biolegend; 405307) secondary antibody was used. The percent of tCD19^+^, KRT5^+^, and P63^+^ cells was quantified using a BD Pharmingen Fortessa-5 laser flow cytometer. Cells stained with isotype control served as a negative control for KRT5 and P63. Unedited cells served as the negative control for tCD19. Flow cytometry data was analyzed using FlowJo (BD Biosciences)

#### Measuring allelic frequency using ddPCR

Gene insertion was validated and allelic frequency was determined via ddPCR. Cells were suspended using TryPLE and washed once with Opti-MEM. 50-100,000 cells were pelleted and resuspended in 50µL of Quick Extract (QE; Lucigen; QE09050) to extract gDNA. The QE suspension was heated at 65°C for 6min and then 98°C for 10min to inactivate QE. The modified *CFTR* locus was amplified using the primers forward: 5′-AGCATCACAAATTTCACAAATAAAGCA-3′ and reverse: 5′-ACCCCAAAATTTTTGTTGGCTGA-3′; and amplification was quantified using the probe sequence: 5′-CACTGCATTCTAGTTGTGGTTTGTCCA-3′. A region in intron 1 of *CFTR* was used as a reference for allele quantification using ddPCR. The reference region was amplified using the primers forward: 5′-TGCTATGCCAGTACAAACCCA-3′ and reverse: 5′-GGAAACCATACTTCAGGAGCTG-3′; and amplification was quantified using the probe sequence: 5′-TTGTTTTTGTATCTCCACCCTG-3′. The PCR reaction for the amplification of the target and reference sequences were as follows: (1) initial denaturation at 95°C for 10min, (2) denaturation at 94°C for 30sec, (3) primer annealing at 56.1°C for 30sec, (4) extension at 72°C for 2min, repeat steps 2-4, 50 times and then (5) infinitely hold at 12°C. The ramp rate for each step was 2°C/s. The samples were then analyzed using the BioRad QX Droplet Reader.

#### Annexin V and propidium iodide staining

Non-CF ABCs were genome edited using the universal strategy. Annexin V apoptosis detection kit with propidium iodide (PI) from Biolegend (Catalog: 640914) was used as per the instruction manual. Briefly, cells were washed twice with cold BioLegend’s Cell Staining Buffer (Catalog: 420201), and then resuspend cells in 100 µL of Annexin V Binding Buffer. 5 µL of FITC Annexin V and 10 µL of PI were added. The cells were gently vortexed and incubated for 15 min at room temperature (25°C) in the dark. 400 µL of Annexin V binding buffer was added to each tube. ABCs treated with only FITC Annexin and only PI were used to set compensation parameters. Annexin V and PI staining in unedited control ABCs and ABCs edited in the presence of AZD7648 and/or ART558 were then evaluated.

#### Off-target Next-Generation Sequencing

After passaging and expanding corrected cells for 4-5 days, they were suspended in TryPLE and washed with Opti-MEM. Cells (50-100,000 cells) were pelleted and resuspended in 50µL of Quick extract (QE; Lucigen; QE09050) to extract gDNA. The QE suspension was heated at 65°C for 6min and then 98°C for 10min to inactivate QE. The one known off-target site was amplified using the primers forward: 5’-ACACTCTTTCCCTACACGACGCTCTTCCGATCTCGACATCCCTTCCCTGAGCCTCTCT-3’ and reverse 5’-GACTGGAGTTCAGACGTGTGCTCTTCCGATCTCTGCACTCCAGCCTGAGCAA-3’.

The DNA was purified using the GeneJet Gel Extraction Kit (ThermoFisher; K0691) according to the manufacturers protocol and sent to Azenta for NGS. The percent of modified alleles different from the reference sequence was quantified using Azenta’s pipeline.

#### ALI culture of corrected ABCs

Cells were suspended using TryPLE 4-5 days after passaging or MACS enrichment. Cells (30-60,000 cells) were plated in 6.5mm Transwell plates with a 0.4µm pore polyester membrane insert (STEMCELL Technologies; 38024). PneumaCult-Ex Plus (500µL) and 0.5µL of 10µM rock inhibitor was added to the basal compartment and 200µL PneumaCult-Ex Plus, 0.2µL of 0.5µM rock inhibitor, and 0.5µL of iMatrix were added to the apical compartment of each well. The cells remained in this medium until reaching confluence at which time the apical medium was removed and the basal medium was replaced with ALI medium obtained from the University of North Carolina Cell Core. The cells were allowed to differentiate for 4-weeks.

#### Ussing chamber functional assays

Ussing chamber measurements were performed on edited CF donor and non-CF donor cells differentiated on ALI transwells for a duration of 4 weeks. For chloride-secretion experiments solutions were as follows: apical: 120mM Na(gluconate), 25mM NaHCO_3_, 3.3mM KH_2_PO_4_, 0.8mM K_2_HPO_4_, 4mM Ca(gluconate)_2_, 1.2mM Mg(gluconate)_2_, and 10mM mannitol, and basolateral: 120mM NaCl, 25mM NaHCO_3_, 3.3mM KH_2_PO_4_, 0.8mM K_2_HPO_4_, 1.2mM CaCl_2_, 1.2mM MgCl_2_, and 10mM glucose. The concentration of ion channel activators and inhibitors was as follows: amiloride (10μM, apical), forskolin (10μM, bilateral; Abcam #ab120058), VX-770 (10μM, apical; SelleckChem #S1144), CFTR_inh_-172 (20μM, apical; Sigma Millipore #C2992), and uridine triphosphate (UTP; 100μM, apical; ThermoScientific Chemicals #AAJ63427MC). The Ussing chamber system used was a VCC MC6 (Physiologic Instruments Inc. San Diego, CA) with the U2500 Self-contained Ussing chambers (Warner Instruments, Hamden CT).

#### Trans-electrical epithelial resistance (TEER)

TEER was recorded from the Ussing chamber system prior once currents had stabilized after mounting samples into the U2500 Self-contained Ussing chambers. TEER is expressed in Ohms.cm^2^.

#### Immunoblotting

Cells were lysed in TN1 lysis buffer [50mM Tris pH 8.0, 10mM EDTA, 10mM Na_4_P_2_O_7_, 10mM NaF, 1% Triton-X 100, 125mM NaCl, 10mM Na_3_VO_4_, and a cocktail of protease inhibitors (Roche Applied Science, Indianapolis, IN)]. Immunoblotting was performed as previously described (Wellmerling et al. 2022 FASEB J.). Briefly, 20-40µg of total proteins were separated with SDS-PAGE in 4-15% polyacrylamide gel and then transferred to polyvinylidenedifluoride (PVDF) membranes (Bio-Rad, Hercules, CA). The membranes were blocked with 5% non-fat milk and immunoblotted with primary antibodies against CFTR (769 from the UNC distribution program) or GAPDH (Santa Cruz Biotechnology, Dallas, TX), followed by appropriate HRP-conjugated secondary antibody (Pierce, Rockford, IL). The signals were detected with enhanced chemiluminescence (Clarity Western ECL substrate and Clarity MAX Western ECL substrate, Bio-Rad, Hercules, CA). The protein bands were scanned using ChemiDoc (Bio-Rad, Hercules, CA).

#### Stanford STAMP

CF-donor cells edited using the universal strategy and enriched for tCD19 were expanded for 4-5 days, collected, and the gDNA was extracted using the GeneJET gDNA Purification Kit (Thermofisher; K0722). NGS of each sample was performed at Stanford Molecular Genetic Pathology Clinical Laboratory to quantify mutation in 130 genes (e.g. *TP53, EGFR, KRA5, NRAS*) identified in the Stanford STAMP. ^61–63^ The Stanford STAMP assay has been clinically validated and can detect variants and variant allele fraction as low as 5%.

#### Immunostaining for epithelial cell characterization

CF-donor cells edited in the presence of AZD7648 and differentiated on ALI transwells for a duration of 4 weeks were stained for cytokeratin 5 (KRT5), acetylated tubulin (ACT), and CD66c to identify basal, ciliated, and secretory cells, respectively. The cells were washed with F-12 (1X) Nutrient Mixture (Ham) (Gibco; 11765054) for 10 minutes on ice. The ALIs were fixed in a solution of 3% sucrose and 4% paraformaldehyde in PBS for a duration of 20 minutes on ice and then washed with PBS. Mouse anti-ACT (Sigma-Aldrich; T6793) was diluted 1:8000. 200µL of the diluted anti-ACT antibody was added to the ALI transwells and incubated overnight at 4°C. The primary antibodies were removed, ALI transwells were washed with PBS, and 200 µL of secondary antibody was added to the ALI transwells. AF488 Donkey anti-rabbit (Life Technologies; A21206) was diluted 1:200 in staining buffer. The ALI transwells were incubated in secondary antibody for a duration of 1 hour. The secondary antibodies were removed, washed with PBS, and Recombinant Alexa Fluor 647 anti-Cytokeratin 5 antibody diluted 1:200 (Abcam, ab193895), mouse anti-CD66c conjugated with PE (Thermoscientific; 12-0667-42) and DAPI (Sigma-Aldrich; D9542) diluted 1:1000 in staining buffer were added to the ALI transwells and incubated for 1 hour. The ALI transwells were washed with PBS, placed on slides using Fluoromount-G Mounting Medium (Invitrogen; 00-4958-02), and cover slipped.

#### Microscopy

Transwell slides were imaged on a motorized Eclipse Ti2-E inverted microscope (Nikon Instruments) with a SOLA LED engine (Lumencor) and an ORCA Fusion CMOS camera (Hamamatsu). Semrock filter sets were used for individual DAPI, GFP, TRITC, and Cy5 channels. Multichannel images were captured as Z-stacks using NIS-Elements AR software (Nikon Instruments, v. 5.30) with 20x Plan Apochromat Lambda objectives (Nikon Instruments) for a final resolution of 0.32 µm/pixel. The Z-stacks were compressed into 2D images using the Extended Depth of Focus (EDF) module in NIS-Elements.

#### Statistical analysis

Statistical significance was assessed using Wilcoxon matched-pairs significant rank test (Figure 1). One-way analysis of variance (ANOVA) followed by multiple comparisons using Tukey’s test. Paired T-test (two-tailed) was performed also.

## Acknowledgments

We thank Dr. Matthew Porteus from Stanford University for the plasmids associated with the universal strategy. Research was supported by Cystic Fibrosis Foundation K-Booster (VAIDYA22A0-KB) and NIH (R00 HL151900). The Cure Cystic Fibrosis Columbus (C3) Epithelial Cell Core (ECC) at Nationwide Children’s Hospital (NCH) and The Ohio State University (OSU) provided primary human airway stem cells, advice, and tools for this work, with the help of the NCH Biopathology Center Core and Data Collaboration Team. C3 is supported by a Cystic Fibrosis Foundation Research Development Program Grant (MCCOY17R2), the NCH Division of Pediatric Pulmonary Medicine (MCCOY19Ro), and the OSU Center for Clinical and Translational Science (UL1TR002733).

## Author Contributions

Conceptualization, S.V., J.T.S, R.R, E.C.B; methodology, S.V., J.T.S, R.E.R, E.C.B; investigation, S.V., J.T.S, R.E.R, E.C.B.,K.L.K, C.J.S, S.H.K; R.N and T.A.V. writing, J.T.S, S.V. All authors participated in the design and conception of experiments and provided editorial feedback on the manuscript.

## Declaration of Interests

None of the authors have any conflict to declare.

## Data Availability Statement

The data supporting the findings presented in this study are available on request from the corresponding author, S.V.

## Notes

### Competing Interest Statement

The authors have declared no competing interest.

