## Supplementary Data for "DNA-PKcs Inhibition Improves Sequential Gene Insertion of the Full-Length *CFTR* cDNA in Airway Stem Cells"

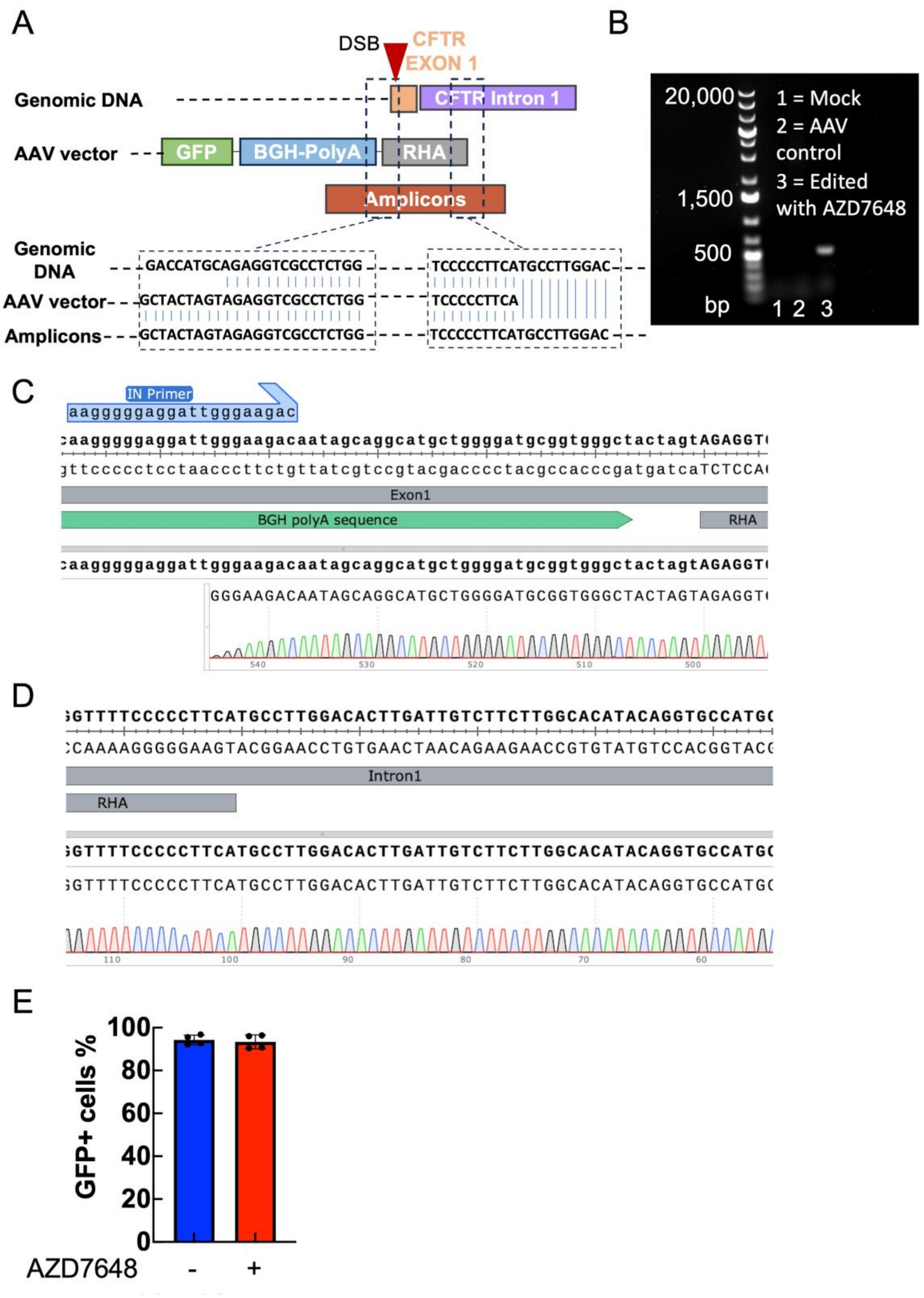


**Figure S1. Insertion of GFP cassette in the CFTR locus.** (A) We used an IN-OUT PCR with primers targeting the polyA tail in the cassette and the genomic DNA in intron 1 of CFTR that was outside the right homology arm (RHA). Amplicons are expected to match the AAV template in the 5' end but only match the genomic DNA at the 3' end. (B) A PCR product was observed only in ABCs edited in the presence of AZD7648 and not in the controls. (C) The sequence of the amplicon corresponds to the polyA tail in the 5' end as expected. (D) The sequence on the 3' end corresponds to the sequence in intron 1 outside the RHA. Thus, the GFP cassette was inserted in the expected locus. (E) Treatment with AZD7648 did not increase AAV uptake in ABCs.

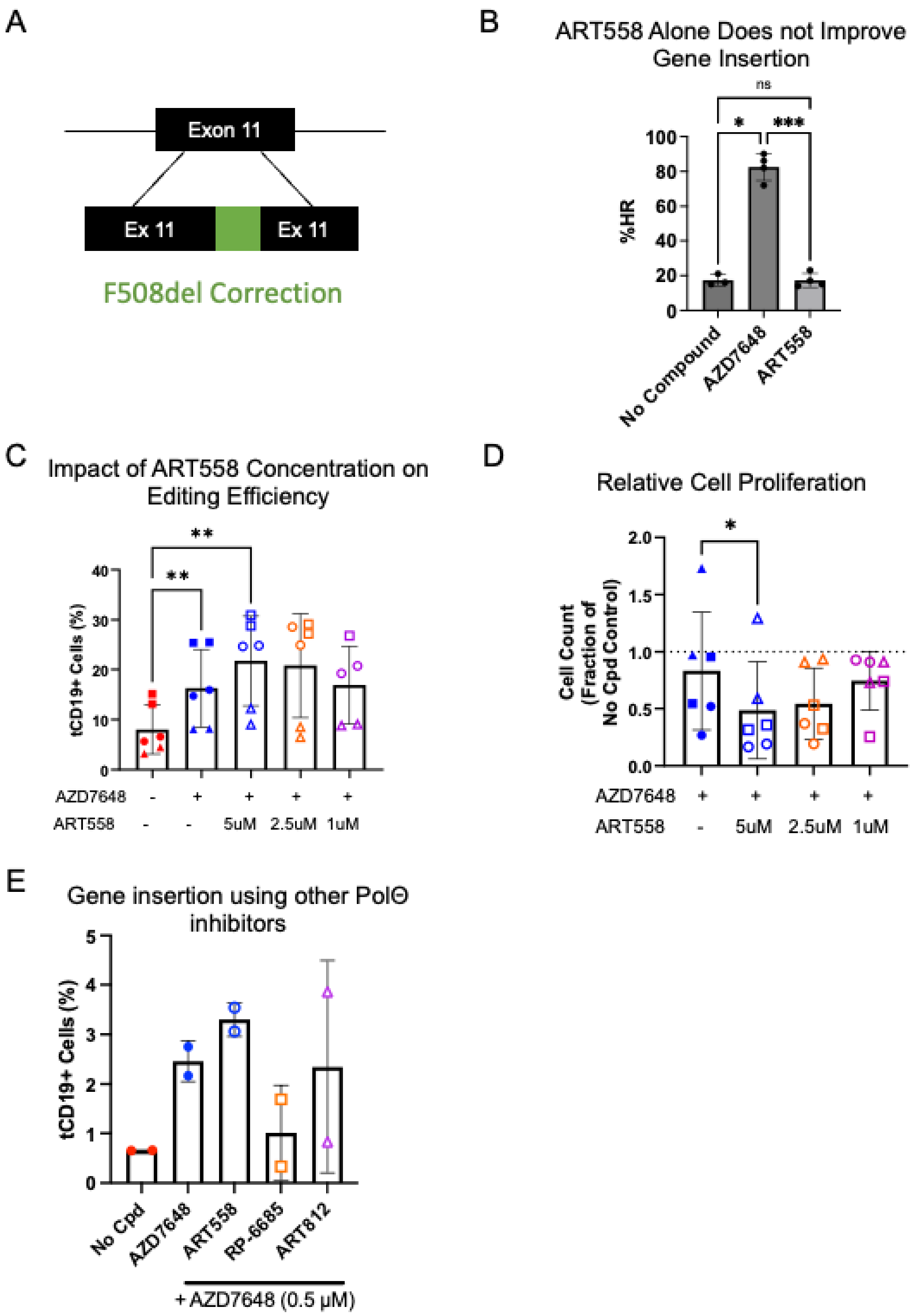


**Figure S2. Impact of ART558 and other PolΘ inhibitors on gene insertion and cell proliferation.** (A) The F508del mutation is present in exon 11 of *CFTR*. We had previously reported the correction of the F508del mutation using CRISPR-Cas9 and AAV.[1] (B) Gene insertion with AZD7648 or ART558 relative to the no compound (n = 2 biological replicates). (C) tCD19 expression in non-CF ABCs in presence of varying concentrations of ART558 (n = 3 biological replicates). (D) Cell proliferation in presence of varying concentrations of ART558 when compared to the AZD7648 only condition (n = 3 biological replicates). (E) tCD19 expression was assessed in non-CF ABCs with the addition of AZD7648 and other PolΘ inhibitors, RP-6685 and ART812, compared to ART558 (n = 1 biological replicate). All statistical analysis was performed with one way ANOVA followed by Tukey's test. ***, **, and * represent p< 0.005, p < 0.01 and p < 0.05 respectively.

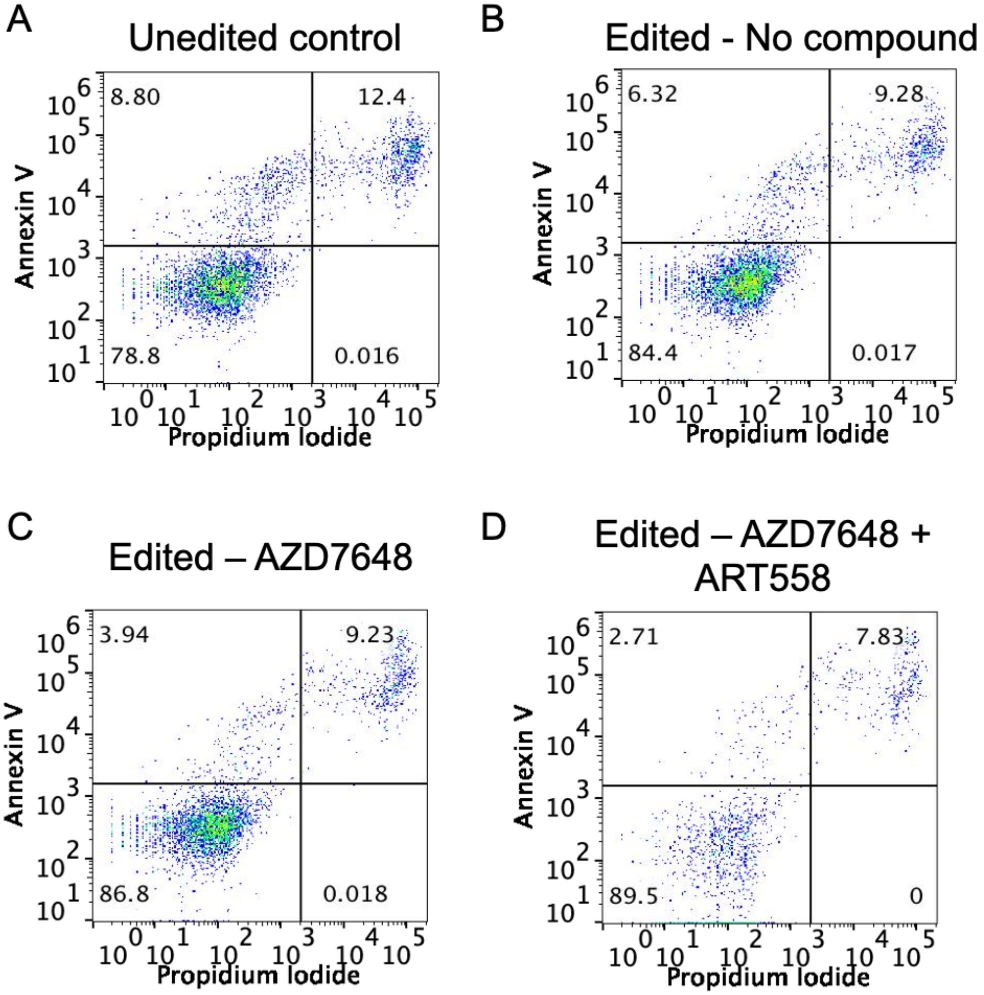


**Figure S3: Annexin V staining and Propidium Iodide Uptake in ABCs Edited in the Presence of AZD7648 and ART558.** ABCs were edited using the universal strategy without the addition of any compounds or with the addition of AZD7648 or both AZD7648 and ART558. The level of editing in the different groups was consistent with the previous results. (A) Unedited ABCs showed slightly more cells positive for Annexin V and propidium iodide than ABCs edited (B) without the addition of any compounds or with the addition of (C) AZD7648 or (D) both AZD7648 and ART558.

**Table S1**: Impact of DNA-PKcs inhibition on CFTR cDNA insertion in ABCs from different donors.

**Universal non-CF donor**

| Donor | %tCD19 | | Fold Change |
|---|---|---|---|
| | **No Compound** | **AZD7648** | |
| 1 | 2.46 | 5.88 | 2.39 |
| 1 | 2.59 | 6.47 | 2.50 |
| 2 | 3.11 | 10.10 | 3.25 |
| 2 | 3.57 | 9.28 | 2.60 |
| 3 | 5.02 | 15.40 | 3.07 |
| 3 | 5.33 | 19.10 | 3.58 |
| 4 | 2.31 | 5.51 | 2.39 |
| 4 | 2.57 | 4.08 | 1.59 |
| 5 | 6.58 | 14.70 | 2.23 |
| 5 | 5.65 | 15.91 | 2.82 |
| 6 | 13.10 | 25.50 | 1.95 |
| 6 | 15.10 | 25.40 | 1.68 |
| 7 | 4.47 | 8.08 | 1.81 |
| 7 | 3.18 | 8.15 | 2.56 |

**Universal CF donor**

| Donor | %tCD19 | | Fold Change |
|---|---|---|---|
| | **No Compound** | **AZD7648** | |
| 8 | 14.80 | 44.90 | 3.03 |
| 8 | 19.00 | 27.90 | 1.47 |
| 9 | 6.42 | 12.80 | 1.99 |
| 10 | 2.88 | 4.09 | 1.42 |
| 11 | 11.90 | 15.30 | 1.29 |
